# Prey’s traits mediate a neotropical toad diet

**DOI:** 10.1101/464511

**Authors:** Matthew T. McElroy, David A. Donoso

**Author notes:** Corresponding author –.

## Abstract

1. Despite the widespread occurrence of myrmecophagy in anurans it is generally unclear whether ant-specialists feed on ants opportunistically or whether they preferentially select for certain species, potentially favoring specific morphological, ecological, or nutritional traits.
2. We flushed 105 stomachs of a lowland neotropical toad, *Rhinella alata*, and identified each consumed ant to species level. We used linear selectivity to calculate predator preference by comparing the abundances of consumed species to their abundances in the leaf litter community on Barro Colorado Island, Panama. We conducted multiple regression models to test whether linear selectivity or general predator preference related to seven morphological characteristics and two measurements of nutritional content.
3. *Rhinella alata* preferentially harvested 24 ant species. Other species were either avoided (n=34) or were eaten opportunistically (n=26). We found that *R. alata* predominantly preys upon large ants that are textured with hair and/or rugosity and preference for prey did not relate to nutrition content. *Rhinella alata* avoided small ants even if they were hyper abundant in the environment, and preferentially ate chemically-defended and aggressive ants if they were large enough.
4. We propose that *R. alata* prefers large ants because they represent a more efficient prey item in terms of predator handling time and because they are easier to see than are smaller ants. Furthermore, we hypothesize that *R. alata* predation attempts are more successful when prey are textured because microstructures on the tongue and prey surface may increase prey adhesion.
5. The ant specialist *R. alata* is not specializing on any particular ant species but rather maximizing prey quantity over quality by only eating the largest ants, despite their scarcity in the environment.

## D. TEXT

Myrmecophagy has evolved multiple times across mammals (Luo, 2017), reptiles (Pianka and Parker, 1975; Pianka and Pianka, 2000), and amphibians (Toft, 1980; Toft, 1981; Simon and Toft, 1991; Daly et al., 1997; Savitsky et al., 2012). The repeated evolution of ant-eating is relatively unsurprising given that ants are conspicuous in both terrestrial and arboreal environments, may constitute 20% of total tropical animal biomass (Fittkau and Klinge 1973), and perform a myriad of ecological and ecosystem functions (Del Toro, Ribbons and Pelini, 2012; Tiede, Donoso, Bendix, Brandl and Farwig, 2017; Roslin et al., 2017). While some predator-ant interactions evolve towards highly specialized interactions (e.g. dart frogs, whose aposematic colors may reflect ant-harvested toxins, (Caldwell, 1996; Santos, Tarvin and O’Connell, 2016), most predators are thought to consume ants opportunistically (Redford, 1986; Griffiths, Greenslade, Miller and Kerle, 1990). However, this general consensus may stem from the lack of diet studies that identify prey taxa to species level (but see Suarez, Richmond and Case, 2000). Though it is well established that ant-eating anurans partition prey based on size (Toft, 1980; Pimentel, 1998; Menendez, 2001), it is unknown if frogs can choose for specific species of ants or for certain ant traits that are available in the surrounding ant community. This uncertainty is particularly evident for tropical anurans whose diets are potentially derived from highly diverse and taxonomically unresolved ant communities.

Tropical leaf litter ant communities are part of complex brown food webs (BFWs, Coleman and Crossley, 2003). BFWs derive their nutrients from detritus (i.e. dead plant material) in a series of little understood processes (Kaspari et al. 2017). Here, most nematodes, mites and collembolans have the task of harvesting wood-decomposing microbes. In turn, these groups feed myriad other insects and invertebrates higher in the food web, such as ants and millipedes. The abundance and diversity of litter critters (that in tropical forests can reach astounding numbers in areas as small as 1m2) is determined by environmental (Donoso and Ramón, 2009; Donoso et al., 2013), biogeochemical (Kaspari et al., 2017), and ecological factors (Moore et al., 2004; Donoso, Johnston and Kaspari, 2010; Donoso, 2017); and are major drivers of ecosystem productivity (Tiede, Donoso, Bendix and Farwig, 2017; Endara et al., 2017; Schuldt et al., 2018). It is in this context that many animals (including frogs) feed upon BFWs’ productivity (Solé, Beckmann, Pelz, Kwet and Engels 2005). However, very few animal ecologists have incorporated BFWs principles into their research. As such, we know little of how BFWs regulate animal biomass (ie. bottom up regulation; Suarez, Richmond, and Case 2000), and in turn how animals control BFW communities (i.e. top-down control).

Very few studies have looked at the specific identities (or traits) of ants in frog diets. This is unfortunate because ants are a morphologically, taxonomically and ecologically diverse group (Del Toro, Ribbons and Pelini, 2012). Therefore, ants provide frogs with potentially different nutrients (Kaspari, Donoso, Lucas, Zumbusch and Kay, 2012), and aposematic alkaloids (Moskowitz et al., In Review; Daly, Garraffo, Spande, Jaramillo and Rand, 1994, Daly, Garraffo, Hall and Cover, 1997, Santos, Tarvin and O’Connell, 2016, Saporito, Spande, Garraffo and Donnelly, 2009). The first of such studies, done in Florida (USA) by Deyrup et al. (2013), found that nocturnal Narrow-Mouthed Toads, *Gastrophryne carolinensis*, fed on up to 43 species of mostly nocturnal ants. These results gave no evidence that frogs present species-level specialization in the ants they eat. Importantly, *G. carolinensis’s* diet included the largest ants in the area (in the genus *Odontomachus* and *Camponotus*, known for their stings and excess formic acid) although in relatively low abundance. More recently, Mcgugan et al. (2016) used mitochondrial COI barcodes to identify ants and mites present in the diets of the dendrobatid frog *Oophaga sylvatica* from three localities in Coastal Ecuador. Mcgugan and collaborators (l.c.) found that alkaloids extracted from the invertebrates differ across geography and, in turn, underlie the alkaloids that are present in *O. sylvatica’s* skin. However, no study has looked at predator preference for specific ants (or traits) by including in the analysis species abundances from the surrounding environment.

*Rhinella alata* is a medium-sized, diurnal, and locally abundant leaf litter toad in Panama. The species is an exceptionally polymorphic and cryptically colored toad (McElroy, 2016), with a diet almost entirely comprised of ants (Toft, 1980; Toft, 1981; Parmelee, 1999; Menendez, 2001; Fajardo, Fajardo and de la Ossa, 2013; Astwood et al., 2016). The toad is chemically defended, but unlike aposematic frogs, predation of ants by *R. alata* should not be driven by toxin sequestration from their diet because *R. alata* synthesizes its own toxins (bufodianaloids) in paratoid glands (Lyttle, Goldstein and Gartz, 1996). Therefore, *R. alata* provides a simplified opportunity to test how prey’s identities, and morphological and nutritional traits, influence predation. Specifically, we answer the following questions, 1) which species of ants does *R. alata* preferentially consume and avoid, and 2) how do prey’s morphological and nutritional traits help to explain its rates of consumption. We conducted this study in the Panama Canal, a tropical seasonal forest that harbors a ~400 species ant community.

## METHODS

### Frog diet collections

Individuals of *R. alata* were sampled from August to September 2010, on Barro Colorado Island (BCI; 09°09’N, 79°51’W) in Panama. BCI receives approximately 2600 mm of annual rainfall, with nearly 90% of it falling between May and November (Leigh, 1999). We opportunistically sampled toads and recorded GPS localities at the point of capture (McElroy, 2015). We recorded Snout-Vent Length (SVL) and removed each individual’s stomach contents using non-lethal gastric lavage (Solé, Beckmann, Pelz, Kwet and Engels, 2005). Stomach contents were stored in 95% ethanol and individual toads were released at the point of capture the following day. We identified individual prey items (i.e. ants) to species-level based on morphology. All other prey items (e.g. mites and Coleoptera) were identified to Class or Family level.

### Sampling the BCI ant community

To sample the ant community within Barro Colorado Island we deployed 216 Berlese funnels at six sites. The sampling was designed and implemented as part of broader studies looking at patterns of morphological trait and phylogenetic dispersion across spatial scales in the island (Donoso, 2014) and the world (Gibb et al., 2017, Parr et al., 2017). More information on methods to survey the leaf litter ant community can be found in Donoso (2014). To reduce noise in our analyses we excluded species that were captured in fewer than five Berlese funnels (leaf litter community dataset) and that were eaten by fewer than two toads (diet dataset). We retained ant species that were eaten frequently but that were not captured in traps, or vice versa, as these potentially represent prey species that *R. alata* highly prefers or avoids. We also excluded army ant species (i.e. *Labidus, Eciton*) from our analysis because *R. alata* specimens, as opposed to army ants, are relatively static, and because army ants can occur in highly temporal high abundance. Thus presence of large numbers of army ants in frog stomachs may not reflect large preference for them. We compared ant abundance and diversity with respect to frog sex and size (SVL) using generalized linear models (GLMs), with a poisson error.

### Prey trait datasets

Our final dataset comprised of 84 ant species and included seven continuous and ordinal morphological traits (weber length; head width; head length; pilosity (i.e. hairy-ness); number of spines; sculpture: smooth - rough; head color: light - dark). Specific information about morphological traits can be found in Table 1 of Parr et al. (2017). To test the hypothesis that prey nutritional content and trophic position are important traits for predator preference we compiled a smaller dataset comprising the 40 species for which nutritional data (percent nitrogen (%N) and isotopic nitrogen (dN)) is available (Kaspari, Donoso, Lucas, Zumbusch and Kay, 2012). Nitrogen composition (high %N = low C:N ratio) is a general proxy for the nutritional value of ants because high %N is positively correlated with protein content, and negatively correlated with Chitin, a molecule indigestible for most vertebrates (Sullivan, Zhang and Bonner, 2013). Stable isotope ratios of nitrogen (dN; expressed here in the per mil ‰ notation) represent a measure of ant trophic position. The abundance of dN (i.e. d15N) of a consumer is typically enriched by 3.4‰ relative to its diet.

**Table 1.** Prey diversity and abundance in toad diets.

|  | Diversity |  |  |  | Abundance |  |  |  |
| --- | --- | --- | --- | --- | --- | --- | --- | --- |
| Diversity | Est. | SE | z | p | Est. | SE | z | p |
| Intercept | 1.22 | 0.77 | 1.58 | 0.11 | 1.30 | 0.31 | 4.24 | <0.001 |
| Sex | -0.64 | 1.40 | -0.46 | 0.65 | -1.05 | 0.55 | -1.90 | 0.06 |
| SVL | 0.01 | 0.02 | 0.74 | 0.46 | 0.05 | 0.01 | 8.44 | <0.001 |
| Sex * SVL | 0.02 | 0.03 | 0.47 | 0.64 | 0.02 | 0.01 | 1.94 | 0.05 |

### Selectivity analysis

To determine *R. alata* preference for, and avoidance of, each ant species we used a linear selectivity index (Strauss, 1979; Palkovacs and Post, 2008; Zandona et al., 2011), which is defined as:

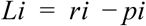

for a prey species i, where Li is the measure of selectivity, ri is the relative abundance in the stomach, and pi is the relative abundance in the environment. To calculate selectivity we pooled the data for stomach contents and subtracted the proportion of a prey species in the environment from the proportion of that prey species in the stomachs. Linear selectivity is easy to interpret; prey species with values > 0 are considered ‘preferred’ and species with values < 0 are considered ‘avoided’. Another benefit of the metric is that it enabled us to retain prey species that were absent from either the stomachs or from the environment. To assess significance of linear selectivity values we replicated our analysis through simulation. We generated 1000 matrices by randomly sampling between 3000-5000 ants (with replacement) from the environmental proportions of ant species. From the simulated matrices we calculated linear selectivity and generated a null distribution of values for each prey species. Then we assigned ant species to three preference categories. We classified ants as preferred if their selectivity values were greater than the null distribution and avoided if their selectivity values were less than the null distribution. We considered species with real selectivity values that fell within the null distribution to be predated upon at a rate that is proportional with their environmental abundance (i.e. ‘neutral’).

### Principal Components Analysis

We characterized the phenotypes of ant species by conducting principal component analysis (PCA) of continuous and categorical trait data. We used the function *dudi.mix* from the package *ade4* because it allows for analysis of both continuous and ordered categorical variables (Dray and Dufour, 2007). The function *dudi.mix* transforms ordinal variables into linear and quadratic variables (i.e. “trait.L” and “trait.Q”, respectively). In the PCA plot the “trait.L” vector points toward increasing values of the trait, while the “trait.Q” vector points in direction of moderate values for the trait. We grouped ant species in the PCA by their preference category to explore which prey traits influence predation.

### Stepwise model selection

We used backwards model selection to determine the best model that explains linear selectivity. We started from a full 5-parameter model and used the function *dropterm* in the packs *MASS* to iteratively remove parameters from our model. We then compared AICc values for each model, calculated Akaike weights to determine our support for each model, and compared coefficients to assess the importance of each trait. Prior to conducting stepwise model selection we used z-scores to standardize the continuous traits. Head width, head length, and weber length were >95% correlated with each other so we only retained weber length as our proxy for size. Because *Ectatomma ruidum* was so highly preferred by frogs (see below) to the point of representing an outlier, we performed a second analysis excluding *E. ruidum* and checked that its inclusion was not skewing the results.

Linear selectivity is a continuous response variable that does not incorporate our null model framework. As such large selectivities values (e.g. *E. ruidum*) are considered outliers and may influence results. To solve this, we performed ordered logistic regression with ant traits as the independent variables and preference category as the response variable. We used backwards model selection and AICc values to determine the best model that explains preference category. We explored the ability of the our model to predict preference category from ant traits. To determine model accuracy, we trained the model with 70% of the data and tested it with the remaining 30%. We bootstrapped the model training 1,000 times and calculated the percent of the time our model predictions matched the testing dataset. When our model predictions did not match the training dataset we distinguished between ‘misses’ (i.e. mis-matches incorporating a neutral preference) and ‘fails’ (i.e. mis-matches between avoid-prefer or prefer-avoid).

### Nutrition Analysis

We analyzed a subset of 40 ant species for which we had %N and dN data to determine whether prey nutrition represents an important but overlooked trait. We conducted ANOVA and Tukey’s posthoc tests to determine whether preferred ant species are larger (WL), have more nitrogen (%N), or are more predatory (dN) than avoided and neutral ant species.

## RESULTS

### Frog diets

We find 3,645 prey items from 61 ant species sampled from 105 individual R. alata. From these, 3,462 items were ants and 183 items (5%)were other taxa. Scarabaeidae (n=47) Curculionidae (n=36) and Aranea (n=34) were the most abundant non-ant taxa. 28 ant species were either army ants or were consumed less than 5 times. Our final dataset thus comprised 3,424 individual ant prey items. The most consumed ant species included *Ectatomma ruidum* (n=2,064), *Pachycondyla harpax* (n=245), *Odontomachus bauri* (n=236), and *Solenopsis* sp. “lash4” (n=213). Across the Winkler trap survey in six sites of the BCI, we collected 26,234 ant specimens from 98 species in 2857 events (Donoso, 2014). The most abundant ants where *Solenopsis* sp. ‘lash4’ (n=4,121), *Wasmannia auropunctata* (n=3,647) and *Solenopsis* ‘JTsp1’ (n=1988). More details of the survey can be found at Donoso (2014). After removing army ants and species too uncommon for analysis, our final community dataset consisted of 25,645 specimens from 84 ant species.

While male frogs were smaller than females (p=<0.001, males: x = 40.6 ± 0.6 mm; females = 46.1 ± 0.9mm), ant diversity in frog stomachs was neither affected by sex (Poisson GLM, Est.=-0.64, p=0.65) nor size (Poisson GLM, Est.=0.01, p=0.46) (Table 1). Instead, larger frogs ate more ants (Poisson GLM, Est.=0.05, p=<0.001) regardless of sex (Poisson GLM, Est.=-1.05, p=0.06) (Table 1)

Ant diversity in frog stomachs was neither affected by sex (Poisson GLM, Est.=-0.64, p=0.65) nor size (Poisson GLM, Est.=0.01, p=0.46). Large frogs ate more ants than small frogs (Poisson GLM, Est.=0.05, p=<0.001). Though females were larger than males (p=<0.001, males: x = 40.6 ± 0.6 mm; females = 46.1 ± 0.9mm) they did not eat significantly more ants than males (Poisson GLM, Est.=-1.05, p=0.06) (Table 1).

### Selectivity analysis

Based on our linear selectivity (Li) analysis and simulations we categorized 23 ant species as preferred, 27 as neutral, and 34 as avoided. *Ectatomma ruidum* was overwhelmingly the most preferred ant species (Li = 0.60), while *Pachycondyla harpax* (Li = 0.07), *Odontomachus bauri* (Li = 0.07), and *Solenopsis* sp. “lash4” (Li = 0.06) represented a second tier of preferred prey items (Figure 1). The first and second most avoided ant species were *Solenopsis* sp. “yellow” (Li = -0.16) and *Wasmannia auropunctata* (Li = -0.13) (Figure 1).

**Figure 1.**
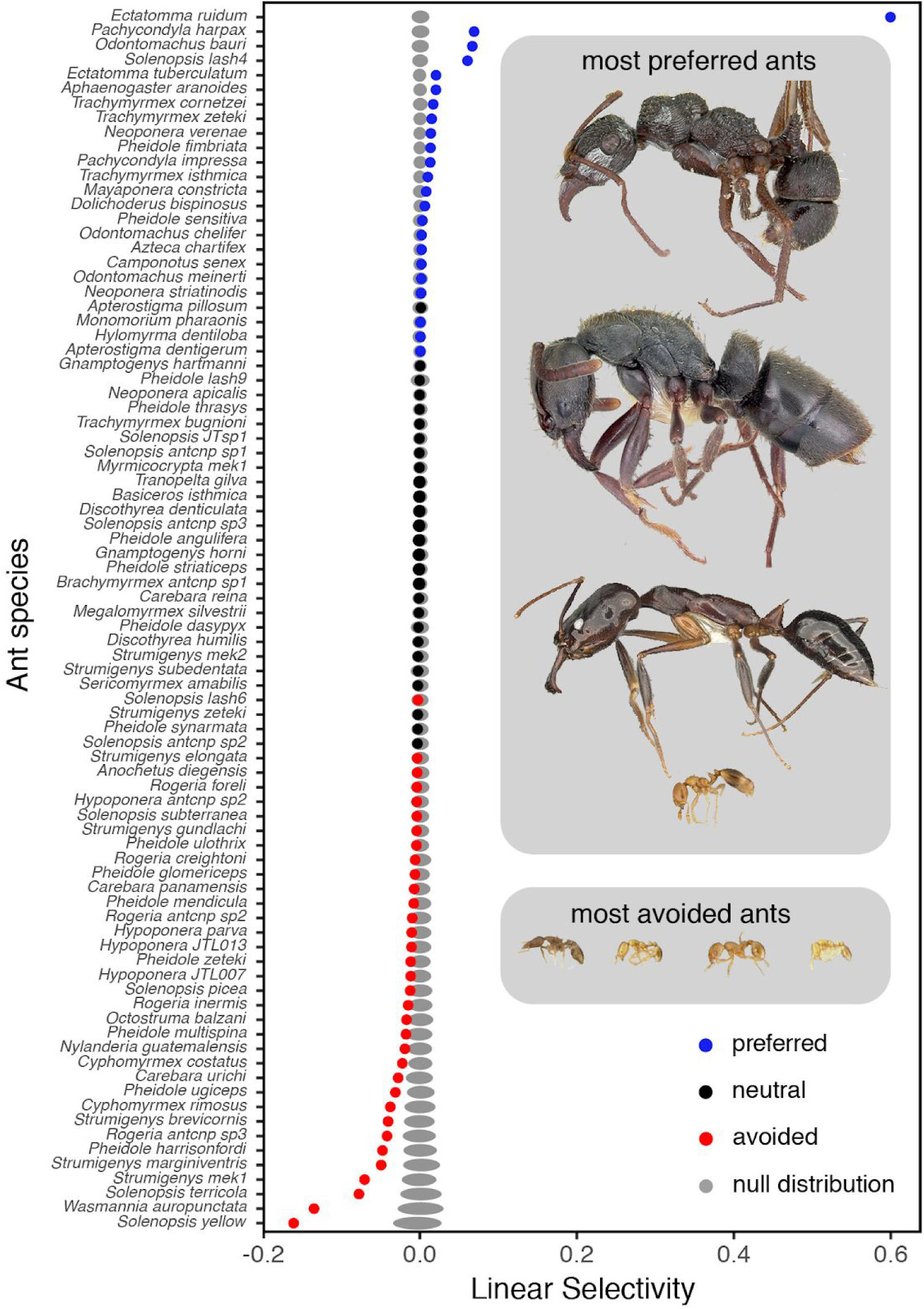
Linear selectivity values for 84 ant species. Preference categories of ant species [prefered (blue), neutral (black) and avoided (red)] were established according to a null model distribution (grey area). Photographs of the top four and bottom four ants are size-scaled in relation to each other.

### Principal Components Analysis

The first principal component accounts for 38.4% of the variation and represents size traits (i.e. weber length, head length, and head width), pilosity, and headcolor.L. The second principal component accounts for 16.8% of the variation and is primarily characterized by traits relating to texture (i.e. sculpture.L, spines, and headcolor.Q). After overlaying the preference categories on our PCA, the results show that neutral and avoided ants largely overlap in trait space, but preferred ant species are larger, darker, and hairier (Figure 2).

**Figure 2.**
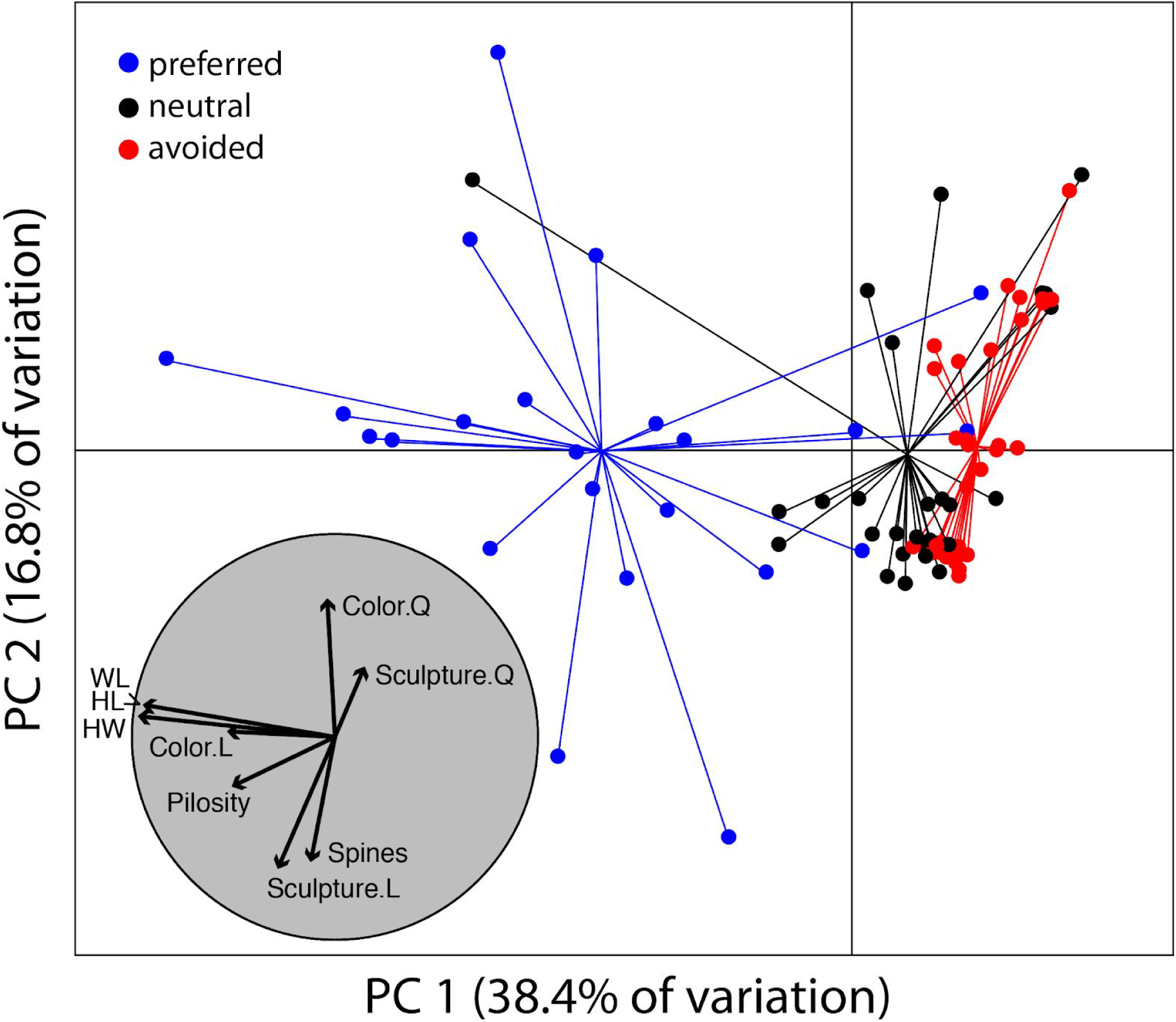
Principal Component Analysis of seven ant traits. Ant species were plotted in the PCA according to preference categories [prefered (blue), neutral (black) and avoided (red)]. Trait.L and Trait.Q are linear and quadratic representations of ordinal variables, respectively.

### Model Selection

The AIC model ‘weber length + pilosity’ was almost three times more informative than the second best model (‘weber length + pilosity + sculpture’) at explaining linear selectivity (Akaike weights = 0.57 vs. 0.21). We found a steep drop off in the ability of subsequent models to explain linear selectivity (Table 2). Removing *E. ruidum* from the analysis did not influence the top models. However, the one parameter model (just ‘pilosity’) dropped from 3rd (Akaike weight = 0.09; Table SM1) to 5th (Akaike weight = 0.01) likely because *E. ruidum* is very pilose. The model rankings for the ordinal logistic regression analysis were consistent with the linear regression analyses. The top three models explaining preference category (Table 3) and linear selectivity (Appendix S1; excluding *E. ruidum*) include the ant traits weber length, pilosity, and sculpture, and have combined probabilities >90%. We utilized the second best model (‘weber length + pilosity + sculpture’) for predicting preference category from ant traits because it includes all the variables in the 90% confidence set. The chosen model correctly predicted the preference category in our training set 70% of the time (x = 0.70; s.d. = 0.10; Figure S1A). When the model incorrectly predicted preference category it was due to ‘misses’ 93% of the time (x = 0.93; s.d. = 0.10; Figure S1B) and ‘fails’ only 7% of the time (x = 0.07; s.d. = 0.10; Figure S1C). Moderately-sized ants that were textured were more likely to be preferred, and small ants that were textured were more likely to be neutral (Figure S2).

**Table 2.** Ranked models (linear response = Linear Selectivity).

| **Model** | **K** | **logLik** | **AICc** | **Delta** | **W** |
| --- | --- | --- | --- | --- | --- |
| Weber Length + Pilosity | 2 | 110.3 | -212.2 | 0.00 | 0.57 |
| Weber Length + Pilosity + Sculpture | 3 | 111.6 | -210.1 | 2.02 | 0.21 |
| Pilosity | 1 | 107.3 | -208.3 | 3.81 | 0.09 |
| Weber Length | 1 | 107.1 | -208.0 | 4.17 | 0.07 |
| Weber Length + Pilosity + Sculpture + Spines | 4 | 111.6 | -207.8 | 4.40 | 0.06 |
| Weber Length + Pilosity + Sculpture + Spines + Color | 5 | 111.8 | -195.3 | 16.83 | <0.001 |

**Table 3.** Ranked models (ordinal response = Toad Preference).

| **Model** | **K** | **logLik** | **AICc** | **Delta** | **weight** |
| --- | --- | --- | --- | --- | --- |
| Weber Length + Pilosity | 2 | -57.8 | 124.0 | 0.00 | 0.61 |
| Weber Length + Pilosity + Sculpture | 3 | -56.6 | 126.3 | 2.26 | 0.20 |
| Weber Length | 1 | -60.4 | -127.1 | 3.05 | 0.13 |
| Weber Length + Pilosity + Sculpture + Spines | 4 | -56.5 | 128.4 | 4.42 | 0.06 |
| Weber Length + Pilosity + Sculpture + Spines + Color | 5 | -54.5 | 137.5 | 13.47 | <0.001 |
| Pilosity | 1 | -78.5 | 163.3 | 39.25 | <0.001 |

### Nutrition

Weber length differed between preference categories (ANOVA: p < 0.001). Preferred ants were two and a half standard deviations larger than avoided ants (TukeyHSD: p < 0.001) and two standard deviations larger than neutral ants (TukeyHSD: p < 0.001). Neutral ants were slightly larger than avoided ants (TukeyHSD: p = 0.03). Neither %N (ANOVA: p = 0.97) nor dN (ANOVA: p = 0.37) differed between preference categories (Figure 3).

**Figure 3.**
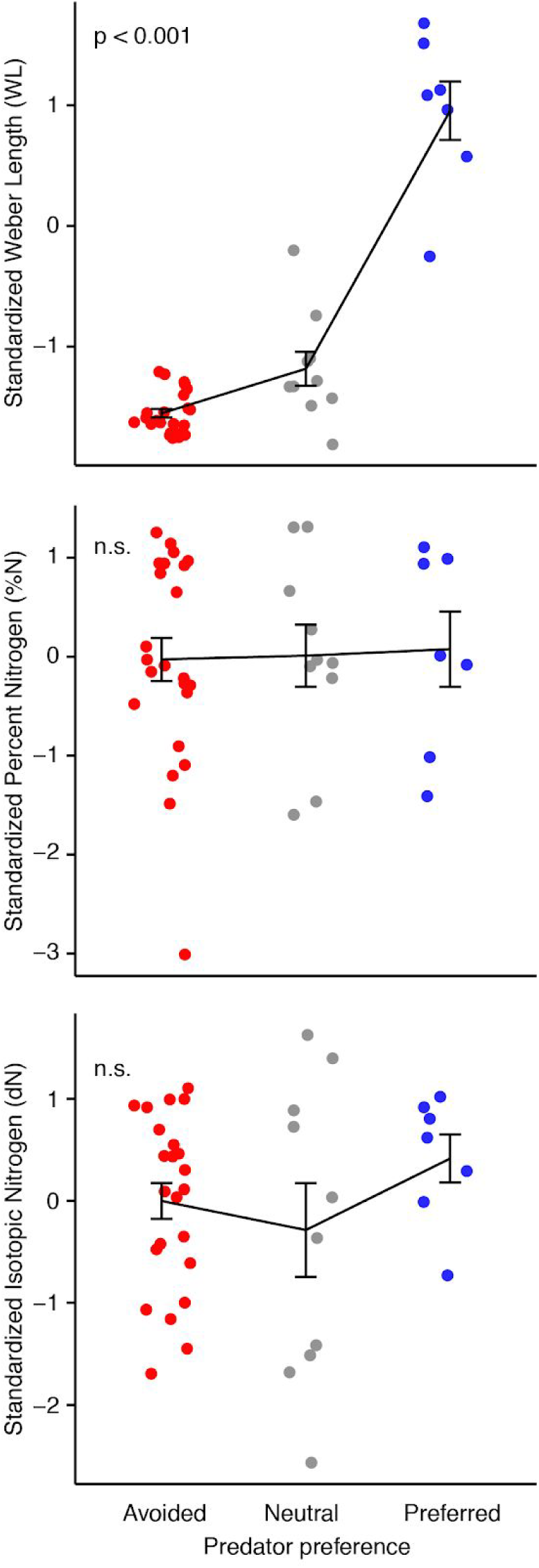
Mean size and nutrition trait values for preferred, neutral, and avoided ant species. Error bars represent 1 standard error.

## DISCUSSION

This is the first study to compare the ant species in frog diets with those in the surrounding leaf litter environments for a diverse tropical forest ecosystem. While it was well established that *Rhinella alata* eats ants, previous studies did not identify prey to species level and they did not compare consumed prey to their availability in the community. Our study found that *R. alata* prefers to eat rarer large ants over the more common smaller ants. Furthermore, we found no relationship between predator preference and nutrition content of prey. Thus, *R. alata* does not search for individual prey items that are high in nutritional value. Instead they secure nutrition by foraging for large ants that provide a larger net nutritional gain, a pattern generally expected from foraging theory.

The best models explaining selectivity and preference include elements of texture, namely hairiness (i.e. pilosity) and rugosity (i.e. sculpture). This is the first field study to find that prey traits other than size can explain predation rates. However, it is difficult to envision a scenario in which *R. alata* is able to discern between - and differentially prey upon - textured and untextured ants. Instead, it is likely that *R. alata* preferentially selects for large ants and that predation success increases with increasing prey texture. Frog tongues are highly specialized for fast and reliable adhesion to prey items. Salivary glands located within the tongue increase the production and availability of highly viscous saliva and are instrumental in the tongues ability to adhere to prey items (Kleintech and Gorb, 2015a; Noel and Hu, 2018). Microstructures on the tongue are thought to increase the adaptability of the tongue to attach to uneven prey surfaces and facilitate the development of mucus fibrils which sustain prey adhesion during tongue retraction (Kleintech and Gorb, 2015b; Kleintech and Gorb, 2016). How variable prey texture (e.g. fur, hair, feathers) may influence tongue adhesion remains poorly understood (Kleintech and Gorb, 2015a). However, microstructures of prey (i.e. pilosity and rugosity) may be analogous to microstructures on the tongue by promoting mucus fibrils and increasing the strength or length of tongue adhesion. Thus, we hypothesize that adhesion between an *R. alata* tongue and a prey item is greater for textured ants than it is for untextured ants, which increases the probability of successfully handling and consuming pilose and rugous ant species. Understanding more about the general functional benefits of texture traits (e.g. pilosity and sensory capability; sculpture and desiccation resistance) and the potential trade-offs associated with predation deserves investigation.

*Ectatomma ruidum* is by far the most selected species in our study, representing 60% of the stomach sample but < 0.01% of our environmental sample. *Ectatomma ruidum*, a common item in Neotropical frog diet studies (Weber, 1938; Lopez, Ghirardi, Scarabotti and Medrano, 2007), has all of the characteristics we expect in a preferred species: it is one of the largest ants in the island, and it is among the most textured species in our dataset. Even though %N and dN are not good predictors of *R. alata* preference, it is intriguing that *E. ruidum* is one of the most nutritious ant species in terms of percent nitrogen (4th of 40) and trophic position (7th of 40). Importantly, size and pilosity predict *R. alata* predation even when *E. ruidum* is removed from the analyses, or when we analyze preference categories (as opposed to linear selectivity). Therefore, the extraordinary presence of *E. ruidum* in *R. alata*’s diet provides a natural validation of our result.

*Wasmannia auropunctata* and other weedy ants in the genus *Solenopsis* were the least prefered. While we know little about these species biology (e.g. we ignore specific chemical or physical defenses these ants may have), *W. auropunctata* is known to displace other animals in Africa (Arnan et al., 2018) and *Solenopsis* include the fire ants with powerful stings. Nonetheless, Solenopsis ants are the prefered food items for the poison frog *Oophaga histrionica* in Colombia (Osorio, Valenzuela, Bermúdez and Castaño, 2015), and in our analysis chemically-defended *Solenopsis* could be preferred if they were large enough (i.e. *Solenopsis* sp. “lash4”). Though we did formally test for avoidance of defensive traits in our analysis, we were unable to identify any patterns relating predator avoidance to other hypothesized defensive traits. For example, we find Army Ants (Dorylinae) in stomachs indicating that aggressive ants with pincers were not necessarily a predator deterrent. Furthermore, *R. alata* preferred large Ponerine ants (e.g. *Pachycondyla harpax* and *Odontomachus bauri*) known for their painful stings (see Deyrup, Deyrup, and Carrel 2013) and they preferred *Trachymyrmex* ants (e.g. *T. isthmica* and *T. zeteki*) that had the most spines (Parr et al. 2017). Taken together, these results indicate that ant traits that are traditionally thought of as anti-predator defenses are unlikely to play a large role in deterring ant-specialist predators. Instead, prey size and foraging behavior likely dictate predation moreso than do anti-predator defenses.

That *R. alata* are eating the largest (and most rare) ants is concerning given that the largest ants are vulnerable to global climate change (Gibb et al. 2015; Gibb et al., 2018) and bottom-up trophic cascades can impact forest food webs (Lister and Garcia, 2018). Furthermore, this study compliments previous work showing that ant communities on BCI are structured by sets of phylogenetically similar ants of small size (Donoso 2014). The preferential predation of large ant species may be partially responsible for producing this pattern (Abrams and Rowe 1996; Roslin et al. 2017). It is difficult to draw general conclusions on the impact that prey traits have on myrmecophagous predator preference until more species-level diet studies accumulate. We provide one example for a non alkaloid-sequestering myrmecophagous frog that clearly selects for large ants despite their scarcity in the environment. Our intriguing finding that pilosity and rugosity influence predation highlights that prey texture may be an overlooked factor in studies on the biomechanics of prey-capture.

## E. AUTHORS’ CONTRIBUTIONS

MTM and DAD conceived the ideas, designed methodology and collected the data; MTM analyzed the data; DAD led the writing of the manuscript. All authors contributed critically to the manuscript and gave final approval for publication.

## E. ACKNOWLEDGEMENTS

Funding was provided by NSF (DEB 0842038) to Mike Kaspari and Adam Kay and the A. Stanley Rand Fellowship awarded to MTM. We thank J. Shik for sharing hints in ant identification. O. Acevedo, B. Jimenez, and H. Castañeda provided valuable support at BCI. We thank the Wang Lab at UC Berkeley for discussion and suggestions that improved the final manuscript. Research was authorized by ANAM # SEX/AP-3-09 and IACUC #20100430.

## F. DATA ACCESSIBILITY

Data available from the Dryad Digital Repository http://XXXXXXXX

## J. SUPPLEMENTARY INFORMATION

### Appendix S1

**Table S1.** Ranked models (linear response = Linear Selectivity). Without Ectatomma.

| Model | K | logLik | AICc | Delta | W |
| --- | --- | --- | --- | --- | --- |
| Weber Length + Pilosity | 2 | 181.6 | -354.8 | 0.00 | 0.55 |
| Weber Length + Pilosity + Sculpture | 3 | 183.2 | -353.2 | 1.54 | 0.26 |
| Weber Length | 1 | 178.9 | -351.4 | 3.35 | 0.10 |
| Weber Length + Pilosity + Sculpture + Spines | 4 | 183.2 | -350.9 | 3.88 | 0.08 |
| Pilosity | 1 | 176.5 | -346.7 | 8.04 | 0.01 |
| Weber Length + Pilosity + Sculpture + Spines + Color | 5 | 185.3 | -342.2 | 12.58 | <0.01 |

**Figure S1.**
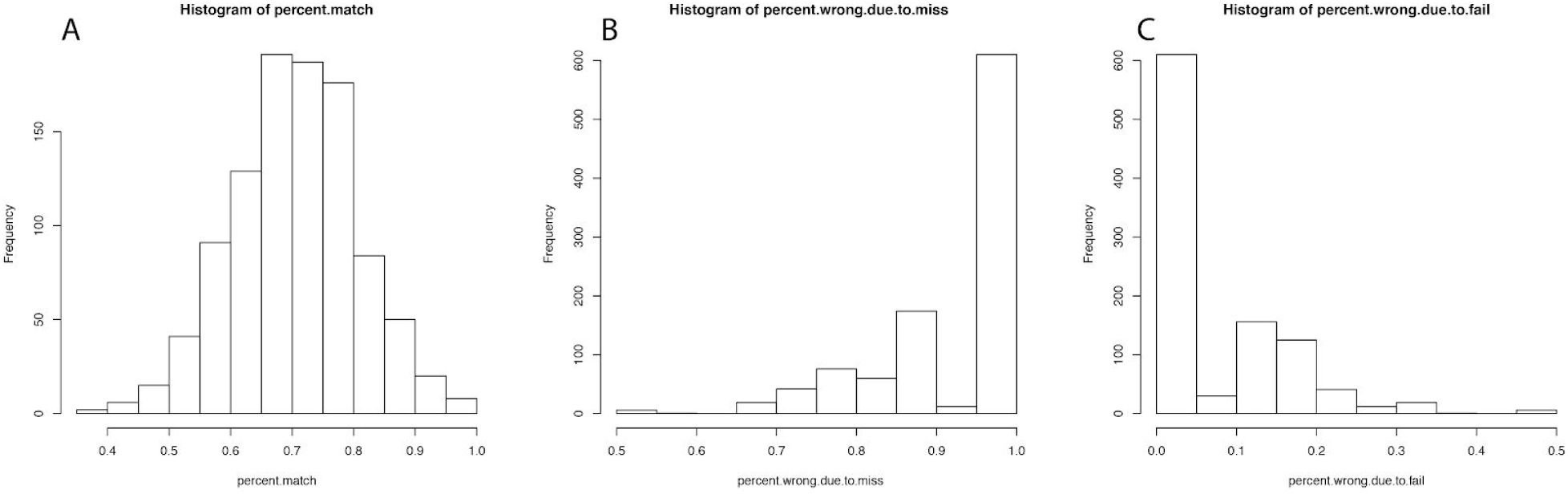
Results for 1000 prediction models for percent match (A), percentage of mis-matches due to “miss” (i.e. involving neutral) (B), and percentage of mis-matches due to “fail” (i.e. predicting avoid instead of prefer, or vice-versa).

**Figure S2.**
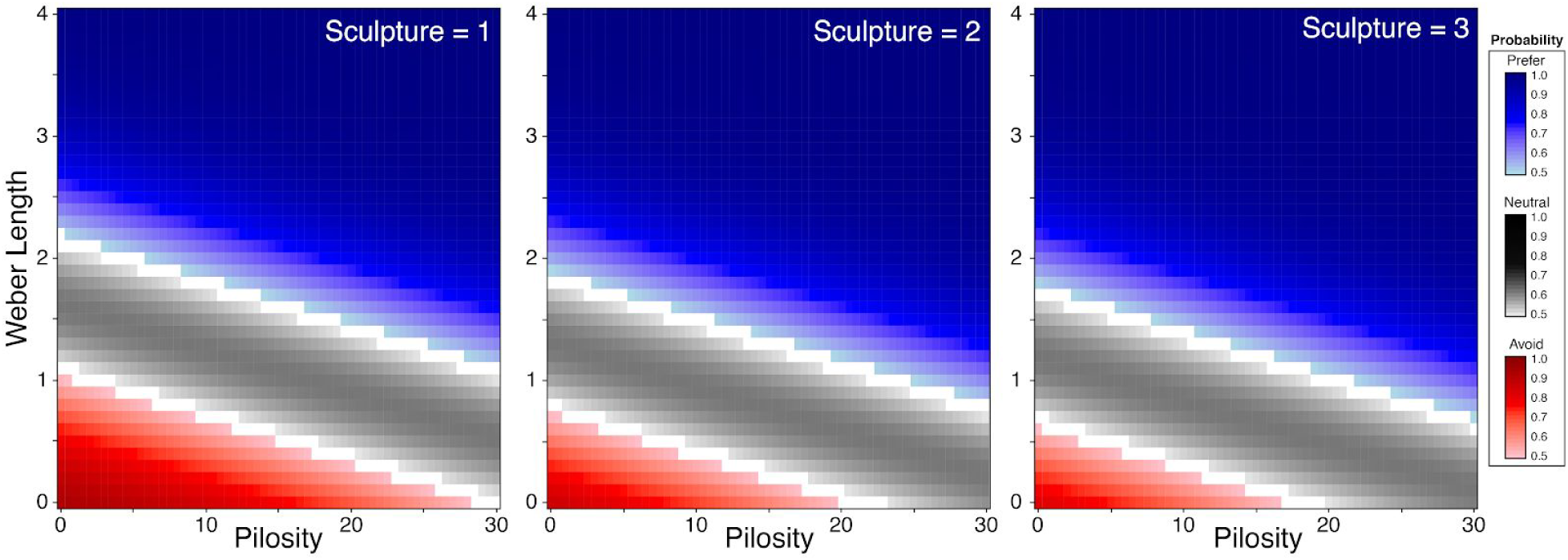
The predictive probability of an ant being preferred (blue), neutral (gray), and avoided based on weber length, pilosity, and sculpture. To aid in visualization we deleted predicted probabilities below 50%.

